# Multistep substrate binding and engagement by the AAA+ ClpXP protease

**DOI:** 10.1101/2020.05.04.076331

**Authors:** Reuben A. Saunders, Benjamin M. Stinson, Tania A. Baker, Robert T. Sauer

**Author notes:** Department of Cellular and Molecular Pharmacology, University of California, San Francisco, California 94158 (Reuben A. Saunders). Biological Chemistry and Molecular Pharmacology, Harvard Medical School, Boston, MA 02115. Correspondence: Robert T. Sauer; MIT 68-533, Cambridge, MA 02139.

## Abstract

*E. coli* ClpXP is one of the most thoroughly studied AAA+ proteases, but relatively little is known about the reactions that allow it to bind and then engage specific protein substrates before the ATP-fueled mechanical unfolding and translocation steps that lead to processive degradation. Here, we employ a fluorescence-quenching assay to study the binding of ssrA-tagged substrates to ClpXP. Polyphasic stopped-flow association and dissociation kinetics support the existence of at least three distinct substrate-bound ClpXP complexes. These kinetic data fit well to a model in which ClpXP and substrate form an initial binding complex, followed by an intermediate complex, and then an engaged complex that is competent for substrate unfolding. The initial association and dissociation steps do not require ATP hydrolysis, but subsequent forward and reverse kinetic steps are accelerated by faster ATP hydrolysis. Our results, together with recent cryo-EM structures of ClpXP bound to substrates, support a model in which the ssrA degron initially binds in the top portion of the axial channel of the ClpX hexamer and then is translocated deeper into the channel in steps that eventually pull the native portion of the substrate against the channel opening. Reversible initial substrate binding allows ClpXP to check potential substrates for degrons, potentially increasing specificity. Subsequent substrate-engagement steps allow ClpXP to grip a wide variety of sequences to ensure efficient unfolding and translocation of almost any native substrate.

**Significance:** AAA+ proteases play key regulatory and quality-control roles in all domains of life. These destructive enzymes recognize damaged, unneeded, or regulatory proteins via specific degrons and unfold them prior to processive degradation. Here, we show that *E. coli* ClpXP, a model AAA+ protease, recognizes ssrA-tagged substrates in a multistep binding and engagement reaction. Together with recent cryo-EM structures, our experiments reveal how ClpXP transitions from a machine that checks potential substrates for appropriate degrons to one that can unfold and translocate almost any protein. Other AAA+ proteases in organelles and bacteria are likely to use similar mechanisms to specifically identify and then destroy their target proteins.

## Introduction

AAA+ proteases play important roles in quality control, protein homeostasis, and cell-cycle regulation by degrading intracellular proteins that are incomplete, damaged, unnecessary, or repress responses to environmental or developmental cues (1, 2). These ATP-fueled proteases consist of a AAA+ ring hexamer and a self-compartmentalized peptidase. Recognition and engagement of the correct protein substrates are the critical initial steps in proteolysis, as subsequent steps have little specificity. In eubacteria and eukaryotic organelles, an unstructured peptide degron at a protein terminus usually targets it for degradation. For the *Escherichia coli* ClpXP protease (3), for example, the AAA+ ClpX ring hexamer binds an unstructured C-terminal or N-terminal degron of a target protein within its axial channel, unfolds any native structure by repeatedly applying force, and then translocates the denatured polypeptide through the channel and into the degradation chamber of ClpP (Fig. 1A). Protein unfolding and translocation by ClpXP have been visualized by single-molecule methods (4–10), but these experiments did not capture the initial steps of substrate binding and engagement, which remain poorly characterized.

**Figure 1.**
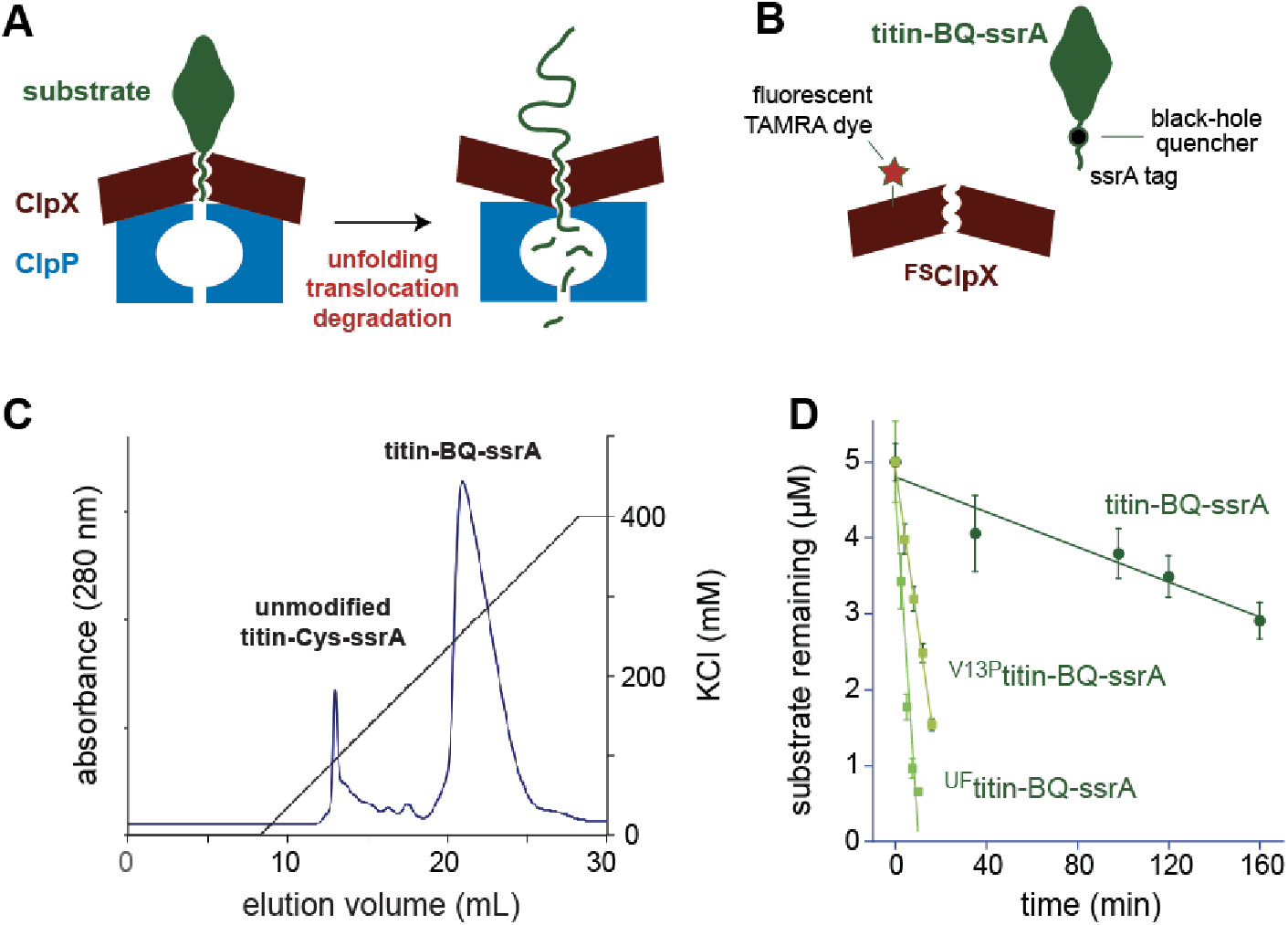
Labeling, purification, and degradation of titin substrates by ClpXP. (A) Cartoon of substrate binding, unfolding, translocation, and degradation by ClpXP. (B) Single-chain ClpX was labeled at residue 170 of one subunit with a fluorescent TAMRA dye, and a cysteine between native titin and the ssrA tag was labeled with a black-hole quencher. (C) Following labeling of the titin-ssrA construct with BHQ-10 maleimide, the modified substrate was separated from unmodified protein by ion-exchange chromatography. (D) Room-temperature degradation of different substrates (5 μM) by ^FS^ClpX (200 nM) and ClpP (600 nM) in the presence of ATP (5 mM) was assayed by SDS-PAGE, staining with Coomassie Blue, and densitometry. Values are averages of two experiments ± SEM. Rates determined from linear fits were 12 ± 0.8 nM min^−1^ (titin-BQ-ssrA), 220 ± 14 nM min^−1^ (^V13P^titin-BQ-ssrA), and 640 ± 40 nM min^−1^ (^UF^titin-BQ-ssrA).

The best-studied degron for *E. coli* ClpXP is the 11-residue ssrA tag, which is cotranslationally appended to the C-terminus of a protein by the ubiquitous tmRNA system when a bacterial ribosome stalls prior to completing synthesis (11, 12). Recombinant addition of this tag to almost any polypeptide or protein makes it a substrate for ClpXP degradation (13–19). ATP or ATPγS must bind to the AAA+ ring of ClpX to support binding to the ssrA tag, but hydrolysis of these nucleotides is not required for ssrA-tag recognition (15, 20–24). During degradation of multidomain substrates, ClpXP often releases a partially degraded product when a very stable native domain is encountered, and single-domain substrates are also likely to be released multiple times prior to degradation (16, 19, 25–28).

Our current understanding of substrate binding and engagement by ClpX or ClpXP is relatively primitive, in part, because previous binding assays have not been compatible with rapid kinetic studies. Here, we describe a robust fluorescence-quenching assay that allows dissection of these important reactions and provide evidence for a model in which an initial binding complex is converted into an intermediate complex and then an engaged complex prior to substrate unfolding and processive translocation/degradation. This model suggests that ClpXP initially checks potential substrates for appropriate degrons and then transitions to a machine that can unfold, translocate, and degrade a wide range of proteins. We discuss these studies in light of recent cryo-EM structures of ClpXP bound to protein substrates (28–30).

## Results

### Binding assayed by fluorescence quenching

To monitor binding of ClpX or ClpXP to an ssrA-tagged substrate by Förster resonance energy transfer (FRET), we labeled ClpX with a fluorescent TAMRA dye and modified an ssrA-tagged substrate containing the I27 domain of human titin by attaching a quencher (Fig. 1B). The ClpX variant was a single-chain enzyme in which each ClpX subunit had its N domain deleted and subunits were linked by short genetically encoded tethers to form a pseudohexamer. We call this variant ^FS^ClpX, where F signifies fluorescent and S indicates single-chain ClpX^ΔN^. Single-chain ClpX pseudohexamers do not dissociate at low concentrations and support degradation of ssrA-tagged substrates by ClpP (31). One subunit of the ClpX pseudohexamer contained a D170C substitution, which represented the only solvent-exposed cysteine and allowed labeling with a fluorescent TAMRA maleimide derivative. The quencher-labeled substrate contained a native titin domain, a C-terminal ssrA tag, and an intervening cysteine to which we attached a black-hole BHQ10 quencher (titin-BQ-ssrA). After labeling with the quencher, we separated the more negatively charged quencher-modified protein from unlabeled protein by anion-exchange chromatography (Fig. 1C). Native titin is a mechanically stable protein that is difficult for ClpX to unfold and degrade (18). We also prepared and purified ^V13P^titin-BQ-ssrA, which is less stable than the parental titin substrate, and ^UF^titin-BQ-ssrA, a variant unfolded by carboxymethylation of otherwise-buried cysteines (18). As assayed by SDS-PAGE, ^FS^ClpXP degraded titin-BQ-ssrA very slowly, ^V13P^titin-BQ-ssrA more rapidly, and ^UF^titin-BQ-ssrA at the fastest rate (Fig. 1D).

We titrated increasing concentrations of titin-BQ-ssrA against a fixed concentration (20 nM) of ^FS^ClpXP (Fig. 2A) or ^FS^ClpX (Fig. 2B) in the presence of saturating ATP or ATPγS, which ClpX or ClpXP hydrolyzes more slowly than ATP (21), determined the degree of quenching after fluorescence had stabilized but before significant degradation could occur, and fit the resulting binding curves to determine apparent affinity constants (*K*_app_). Substrate binding appeared slightly tighter in the presence of ATPγS than ATP and slightly tighter to ClpXP than to ClpX. We detected no binding of ^FS^ClpXP or ^FS^ClpX to titin-BQ-ssrA in the presence of ADP.

**Figure 2.**
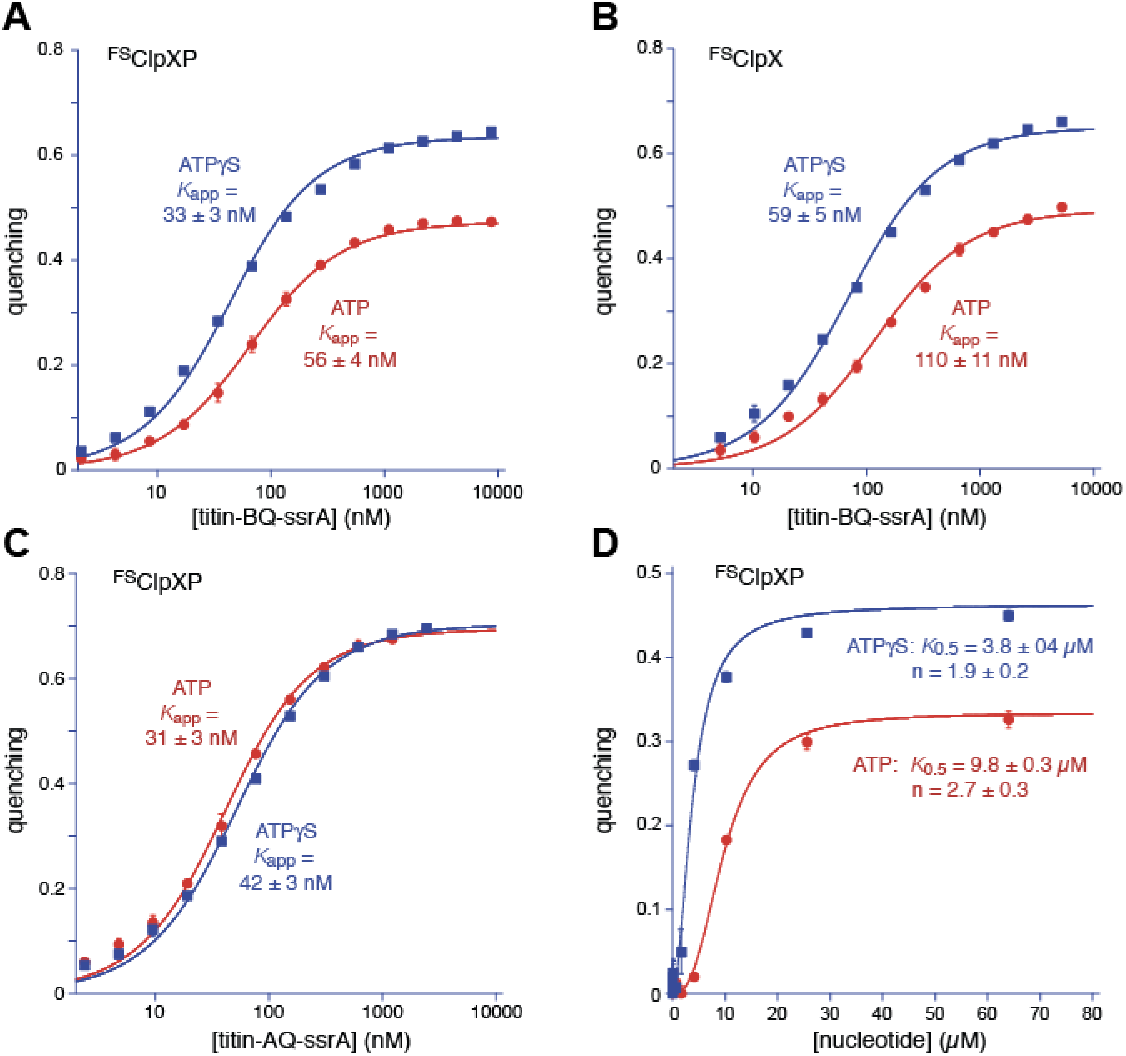
Assays of protein substrate or nucleotide binding to ^FS^ClpXP or ^FS^ClpX. (A) Binding of titin-BQ-ssrA to ^FS^ClpXP (20 nM) in the presence of ATP (5 mM) or ATPγS (1 mM). Values are averages ± SD of three independent experiments and the fits are to a quadratic tight-binding model. (B) Same as panel A but using ^FS^ClpX. (C) Same as panel A but using titin-AQ-ssrA. (D) FS Activation of titin-BQ-ssrA binding to ^FS^ClpXP by addition of ATP or ATPγS. Values are averages ± SD of three experiments and the fits are to the Hill equation (n is the Hill constant).

Higher maximal quenching was observed in ATPγS compared to ATP for both ^FS^ClpXP and ^FS^ClpX (Figs. 2A, 2B). Quenching by FRET is sensitive to small changes in average distances near the Förster radius (32). Thus, relative to ATP, ATPγS may result in ClpX-substrate structures with a closer average distance between the fluorophore and quencher. Alternatively, ATP and ATPγS could result in differences in the fraction of ClpX enzymes capable of binding substrate, even at saturating concentrations. To distinguish between these possibilities, we modulated the quenching Förster radius by labeling the titin substrate with ATTO 575Q (AQ), which has a larger Förster radius when paired with TAMRA (~62 Å) than does BHQ10 (~46 Å). If higher quenching in ATPγS resulted from a higher fraction of binding-competent ClpX enzymes, then higher quenching should also be observed for the titin-AQ-ssrA substrate. However, in titration experiments, titin-AQ-ssrA binding to ^FS^ClpXP resulted in similar fluorescence quenching in the presence of ATP or ATPγS (Fig. 2C). Thus, the ssrA-tagged substrate is likely to be held in slightly different average conformations in the ensembles of ATP-bound and ATPγS-bound ClpXP structures. The conformations of individual states in these ensembles may be identical for both nucleotides, but slower hydrolysis of ATPγS could, for example, result in a higher population of a conformation in which the fluorophore and quencher are closer together. We cannot rigorously exclude the possibility that titin-AQ-ssrA binds somewhat differently to ATP-ClpXP than ATPγS-ClpXP, but this model seems less likely.

Using the quenching assay, we found that ATPγS supported ^FS^ClpXP binding to the titin-BQ-ssrA substrate at lower concentrations than ATP and with modestly reduced positive cooperativity as measured by the Hill constant of the fit (Fig. 2D). Positive cooperativity in the ATP activation of substrate binding to a ClpX mutant defective in ATP hydrolysis was reported previously (23), and likely reflects a requirement that multiple subunits in the ClpX hexamer must bind ATP to stabilize the substrate-binding conformation. In accord with this model, four or five subunits of the ClpX hexamer are ATP or ATPγS bound in substrate-bound cryo-EM structures (28–30). The finding that ATPγS supports substrate binding at lower concentrations and with a lower Hill constant than ATP suggests that faster hydrolysis of ATP at sub-saturating concentrations results in a lower population of nucleotide-bound intermediates that can bind substrate.

### Association kinetics

Stopped-flow experiments were used to study the kinetics of ClpXP binding to substrate. We used a fixed concentration of ^FS^ClpXP and increasing concentrations of titin-BQ-ssrA to initiate binding reactions in the presence of ATP (Fig. 3A). The resulting fluorescence trajectories were polyphasic, indicating that the reaction has multiple steps. At high substrate concentrations, we observed two initial phases of decreased fluorescence, indicating elevated degrees of quenching. Fluorescence then increased as the reaction approached steady state. To test if the rate of nucleoside-triphosphate hydrolysis affected the engagement reaction, we conducted similar experiments in the presence of ATPγS (Fig. 3B). In these experiments, the initial binding phase had kinetics similar to the reaction initiated with ATP. However, the second decreased fluorescence phase was slower and fluorescence recovery also occurred more slowly than in ATP and resulted in a lower steady-state level of fluorescence, consistent with the differences observed in maximal quenching in binding experiments performed with ATP or ATPγS (Fig. 2A).

**Figure 3.**
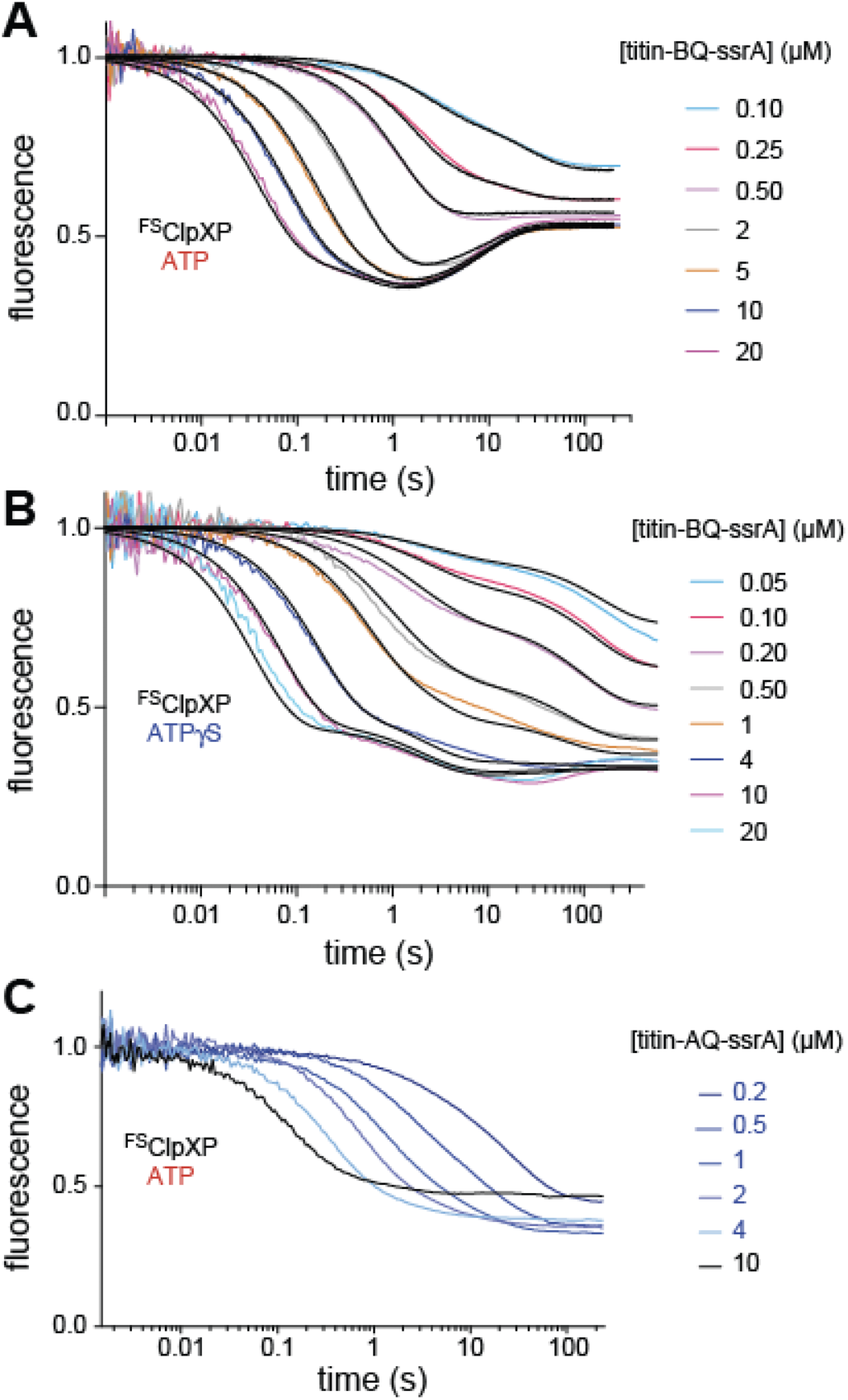
Association kinetics. (A) ^FS^ClpXP (final concentration 10 nM) and ATP (final concentration 2.5 mM) were mixed with the indicated concentrations of titin-BQ-ssrA to initiate binding, which was assayed by changes in fluorescence. (B) Same experiment as in panel A but using ATPγS (final concentration 0.5 mM). (C) Same experiment as in panel A but using the titin-AQ-ssrA substrate. In panels A and B, the black lines represent global fits to the first three steps of the model shown in Figure 5.

We hypothesized that the complex fluorescence trajectories observed in the titin-BQ-ssrA association experiments reflected transitions between free ClpXP and several bound states with different fluorophore-quencher distances and quenching efficiencies, which could represent transient states in the engagement reaction. Alternatively, the fluorescence trajectories could reflect distinct populations of ClpXP that bind and release titin-BQ-ssrA at different rates. To distinguish between these possibilities, we conducted stopped-flow experiments with titin-AQ-ssrA, as TAMRA quenching by AQ should be less sensitive to the fluorophore-quencher distance at relevant length scales, and different bound states are thus more likely to quench with similar efficiencies. Rapid mixing of ^FS^ClpXP and titin-AQ-ssrA caused a monotonic decrease in fluorescence (Fig. 3C), indicating that the more complex fluorescence trajectories observed with titin-BQ-ssrA do not result from distinct ClpXP populations that bind and release substrate at different rates.

### Dissociation kinetics

We conducted stopped-flow dissociation experiments by allowing complexes of titin-BQ-ssrA with ^FS^ClpXP to form in ATP or ATPγS for at least 20 min, adding a large excess of unlabeled titin-ssrA to initiate dissociation, and recording changes in fluorescence (Fig. 4). A biphasic increase in fluorescence was observed in the presence of ATP or ATPγS. In ATP, the fast phase (15% amplitude) had a time constant of ~3 s and the slow phase (85% amplitude) had a time constant of ~50 s. In ATPγS, the fast phase (17% amplitude) had a time constant of ~2 s and the slow phase (83% amplitude) had a time constant of ~240 s.

**Figure 4.**
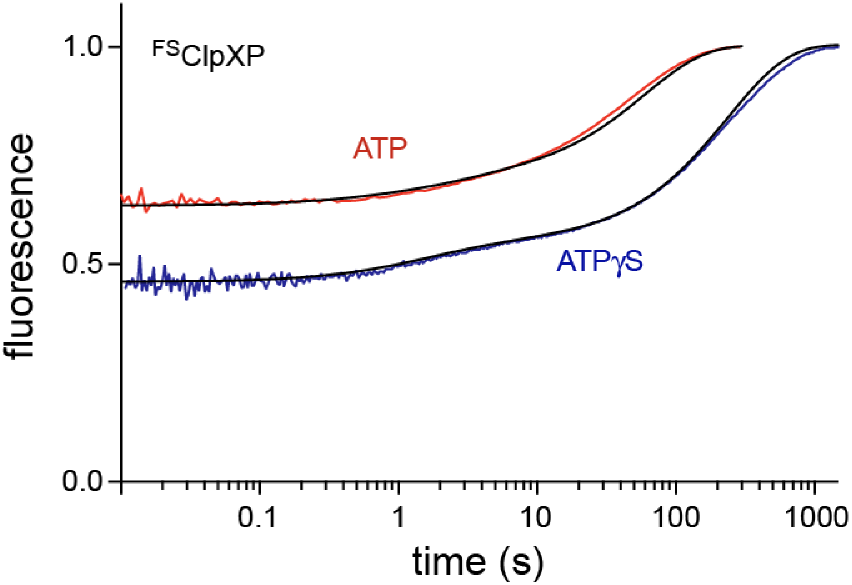
Dissociation kinetics. Complexes of ^FS^ClpXP (20 nM) and titin-BQ-ssrA (1 μM) were allowed to equilibrate in ATP (5 mM) or ATPγS (1 mM), then mixed at time zero with an equal volume of unlabeled titin-ssrA (50 μM) to initiate dissociation, and changes in fluorescence were recorded as a function of time. The black lines are simulations calculated using the first three steps of the Fig. 5 model and the parameters listed in Table 1.

### Fitting to a multistep binding and engagement model

Models containing fewer than three substrate-bound ClpX species did not fit the titin-BQ-ssrA association data and an additional state was necessary to account for translocation and degradation. In the model depicted in Fig. 5, ClpXP associates with an ssrA-tagged substrate (*k*_1_) to form a binding complex (BC). In a subsequent step (*k*_2_), ATP hydrolysis and translocation by ClpXP converts the binding complex into an intermediate complex (IC) in which the ssrA tag moves deeper into the channel, but the native portion of the substrate is not yet in contact with the ClpX ring. Additional translocation (*k*_3_) brings the native portion of substrate in contact with the top of the ClpX ring to form an engaged complex (EC), which can unfold a native substrate. From this state, repeated ATP-fueled power strokes can unfold the substrate resulting in a processive translocation complex (TC) that eventually results in substrate degradation in a multistep process subsumed under the *k*_4_ rate constant. A final step (*k*_5_) clears the axial channel of ClpX, regenerating free enzyme to bind another substrate.

**Figure 5.**
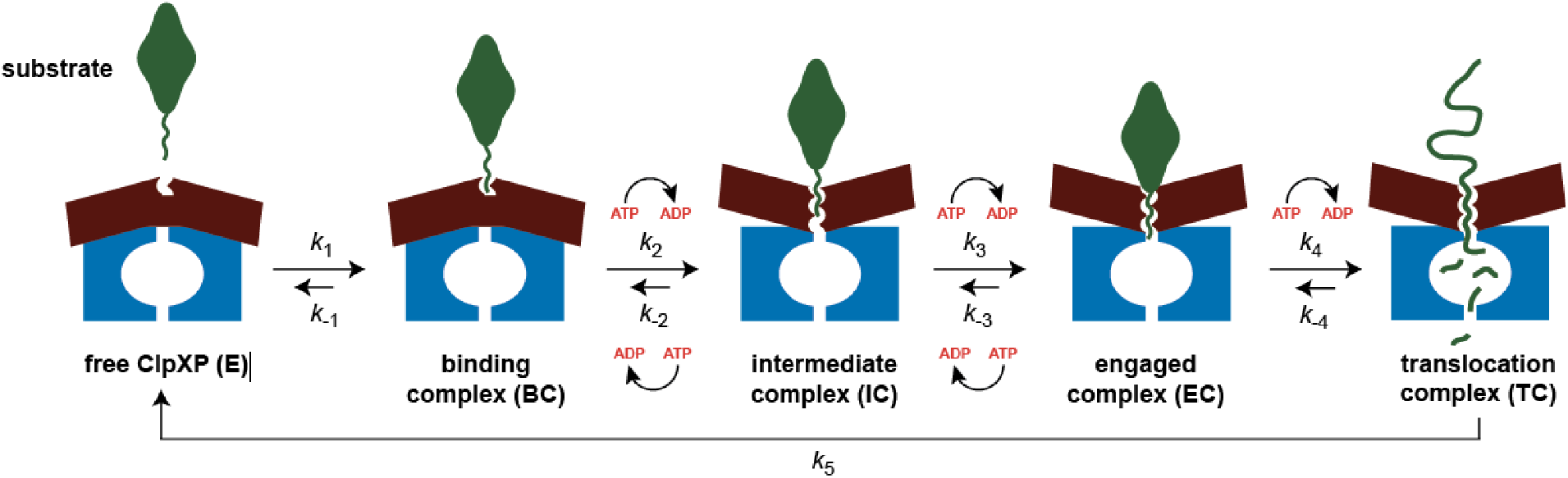
Model for substrate binding, engagement, unfolding, and processive translocation/degradation by ClpXP. As discussed in the text, the *k*_2_, *k*_-2_, *k*_3_, and *k*_-3_ rate constants increase as the rate of nucleoside triphosphate hydrolysis increases. The composite *k*_4_ rate constant for ClpXP substrate unfolding and translocation also depends on the hydrolysis rate (21).

ClpXP unfolding of the titin-BQ-ssrA substrate (included in the *k*_4_ step) is expected to be very slow (18) compared to the earlier binding and engagement steps shown in Fig. 5. Thus, we globally fit the titin-BQ-ssrA association data to determine rate constants for the *k*_1_, *k*_-1_, *k*_2_, *k*_-2_, *k*_3_, and *k*_-3_ steps and quenching values for each state in the presence of ATP or ATPγS (Table 1). As shown in Figs. 3A and 3B, simulations based on the model and extracted rate constants and quenching values provided good fits (black lines) of the association data over a wide range of substrate concentrations. Simulations of dissociation kinetics based on the association model and parameters also resulted in close agreement with the experimental results (Fig. 4).

**Table 1.**
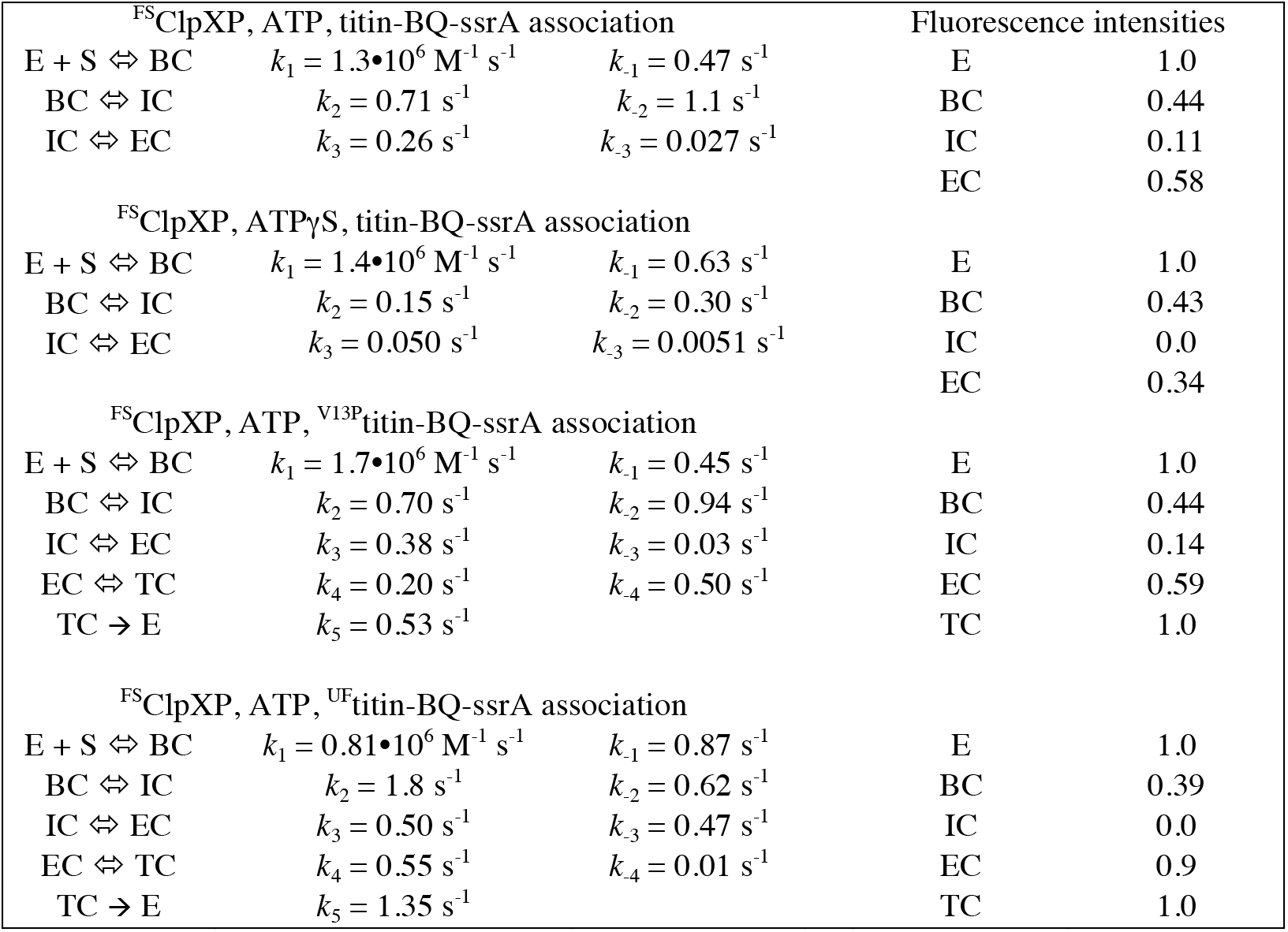
Parameters obtained from global fitting of association experiments using the model shown in Figure 5.

| <sup>FS</sup>ClpXP, ATP, titin-BQ-ssrA association |  |  |  | Fluorescence intensities |  |
| --- | --- | --- | --- | --- | --- |
| E + S ⇌ BC | $k_1 = 1.3 \cdot 10^6 \text{ M}^{-1} \text{ s}^{-1}$ | $k_{-1} = 0.47 \text{ s}^{-1}$ | | E | 1.0 |
| BC ⇌ IC | $k_2 = 0.71 \text{ s}^{-1}$ | $k_{-2} = 1.1 \text{ s}^{-1}$ | | BC | 0.44 |
| IC ⇌ EC | $k_3 = 0.26 \text{ s}^{-1}$ | $k_{-3} = 0.027 \text{ s}^{-1}$ | | IC | 0.11 |
|  |  |  |  | EC | 0.58 |
| <sup>FS</sup>ClpXP, ATPγS, titin-BQ-ssrA association | | | | | |
| E + S ⇌ BC | $k_1 = 1.4 \cdot 10^6 \text{ M}^{-1} \text{ s}^{-1}$ | $k_{-1} = 0.63 \text{ s}^{-1}$ | | E | 1.0 |
| BC ⇌ IC | $k_2 = 0.15 \text{ s}^{-1}$ | $k_{-2} = 0.30 \text{ s}^{-1}$ | | BC | 0.43 |
| IC ⇌ EC | $k_3 = 0.050 \text{ s}^{-1}$ | $k_{-3} = 0.0051 \text{ s}^{-1}$ | | IC | 0.0 |
|  |  |  |  | EC | 0.34 |
| <sup>FS</sup>ClpXP, ATP, <sup>V13P</sup>titin-BQ-ssrA association |  |  |  |  |  |
| E + S ⇌ BC | $k_1 = 1.7 \cdot 10^6 \text{ M}^{-1} \text{ s}^{-1}$ | $k_{-1} = 0.45 \text{ s}^{-1}$ | | E | 1.0 |
| BC ⇌ IC | $k_2 = 0.70 \text{ s}^{-1}$ | $k_{-2} = 0.94 \text{ s}^{-1}$ | | BC | 0.44 |
| IC ⇌ EC | $k_3 = 0.38 \text{ s}^{-1}$ | $k_{-3} = 0.03 \text{ s}^{-1}$ | | IC | 0.14 |
| EC ⇌ TC | $k_4 = 0.20 \text{ s}^{-1}$ | $k_{-4} = 0.50 \text{ s}^{-1}$ | | EC | 0.59 |
| TC → E | $k_5 = 0.53 \text{ s}^{-1}$ | | | TC | 1.0 |
| <sup>FS</sup>ClpXP, ATP, <sup>UF</sup>titin-BQ-ssrA association |  |  |  |  |  |
| E + S ⇌ BC | $k_1 = 0.81 \cdot 10^6 \text{ M}^{-1} \text{ s}^{-1}$ | $k_{-1} = 0.87 \text{ s}^{-1}$ | | E | 1.0 |
| BC ⇌ IC | $k_2 = 1.8 \text{ s}^{-1}$ | $k_{-2} = 0.62 \text{ s}^{-1}$ | | BC | 0.39 |
| IC ⇌ EC | $k_3 = 0.50 \text{ s}^{-1}$ | $k_{-3} = 0.47 \text{ s}^{-1}$ | | IC | 0.0 |
| EC ⇌ TC | $k_4 = 0.55 \text{ s}^{-1}$ | $k_{-4} = 0.01 \text{ s}^{-1}$ | | EC | 0.9 |
| TC → E | $k_5 = 1.35 \text{ s}^{-1}$ | | | TC | 1.0 |

The rate constants for bimolecular association (*k*_1_) of titin-BQ-ssrA and ^FS^ClpXP and dissociation of this complex (*k*_-1_) were similar in the presence of ATP and ATPγS (Table 1), indicating that ATP hydrolysis does not play a substantial role in the initial recognition reaction. For the *k*_2_ and *k*_3_ steps, by contrast, rate constants were 3.5- to 7-fold slower in the presence of ATPγS, suggesting that the rate of hydrolysis is important, as expected if power strokes are required for these steps. Rate constants for the reverse *k*_-2_ and *k*_-3_ steps were also 3.5- to 6-fold slower in ATPγS compared to ATP, indicating that these steps are also hydrolysis dependent and probably involve post-hydrolysis states after one or more unsuccessful power strokes.

### Binding, engagement, and degradation of less stable or unfolded substrates

We also assayed the kinetics of ^FS^ClpXP association with different concentrations of ^V13P^titin-BQ-ssrA, whose native structure is less stable than that of titin-BQ-ssrA, or ^UF^titin-BQ-ssrA, which is unfolded (Figs. 6A, 6B). In both cases, the changes in fluorescence at early times (0 to 1 s) resembled those of titin-BQ-ssrA (Fig. 3A). At later times, differences between the three substrates were most pronounced at low substrate concentrations and could be attributed to faster degradation of the ^V13P^titin-BQ-ssrA and ^UF^titin-BQ-ssrA substrates. Table 1 lists rate constants obtained by fitting the association experiments for ^V13P^titin-BQ-ssrA and ^UF^titin-BQ-ssrA to the full model shown in Fig. 5. The similarity of the early trajectories for the titin substrates, despite large differences in their stabilities and degradation rates, indicates that folding stability does not exert a major influence on the early steps of the binding/engagement process. For the native titin-BQ-ssrA and ^V13P^titin-BQ-ssrA substrates in the presence of ATP, values obtained from fitting for the *k*_1_, *k*_-1_, *k*_2_, *k*_-2_, *k*_3_, and *k*_-3_ rate constants and intensities were roughly similar (Table 1). These values showed more variation for the unfolded titin substrate, including a 16-fold increase in the *k*_-3_ rate constant (Table 1). The presence of native structure in the substrate may influence the kinetics and populations of microstates during the binding and engagement steps, as cryo-EM structures show that substrate-binding loops of ClpX contact native portions of substrates in complexes similar to the engaged complex depicted in Fig. 5 (29). Because unfolding is not necessary for degradation of the ^UF^titin-BQ-ssrA substrate, the EC state and associated rate constants may differ from those of the native substrates. Indeed, the fluorescence intensity of the EC state for the unfolded substrate was substantially higher than those of the native substrates (Table 1), suggesting that the substrate polypeptide may have moved more deeply into the ClpX channel.

**Figure 6.**
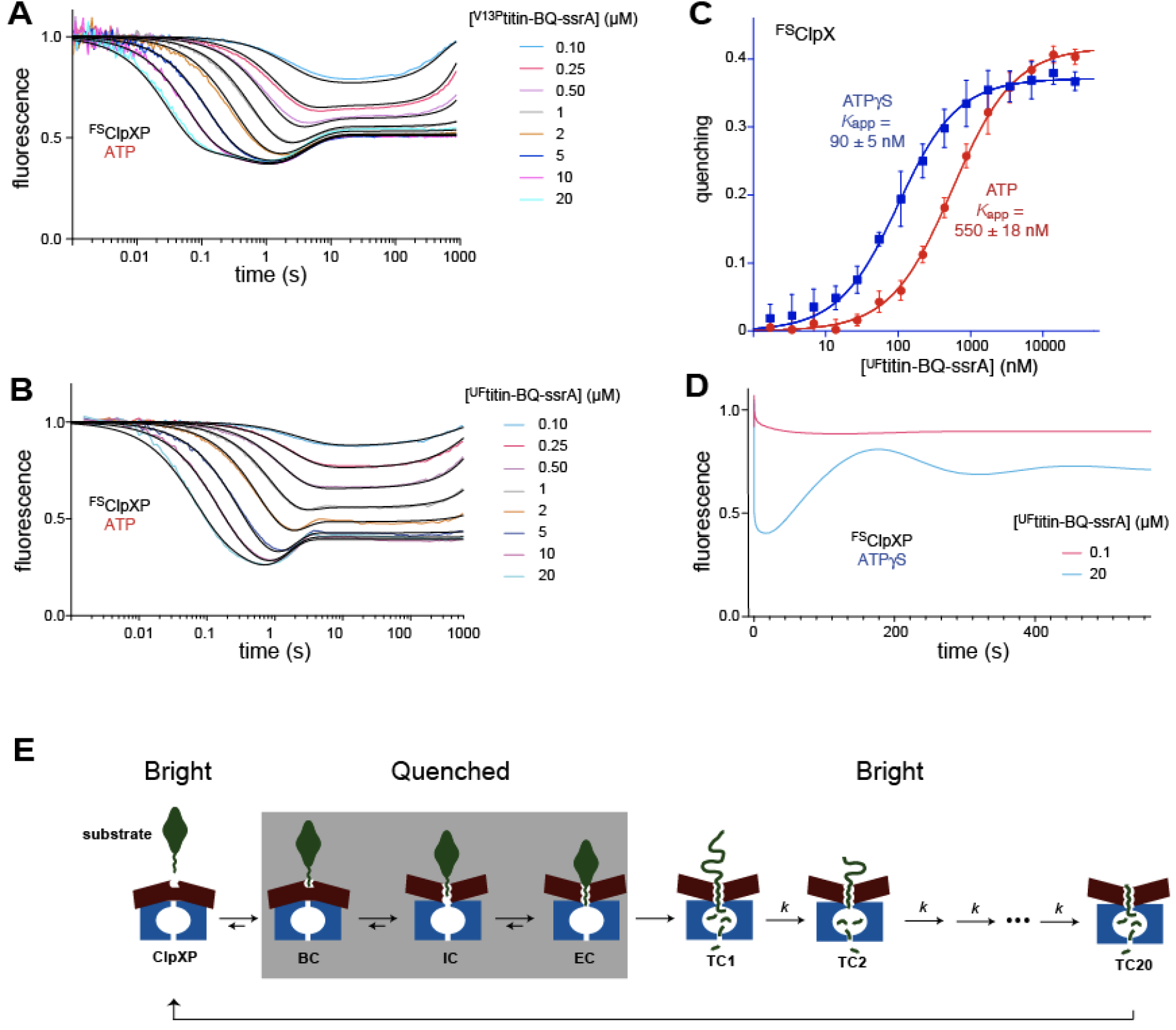
Association, binding, and degradation of less-stable or unfolded titin substrates. (A) ^FS^ClpXP (10 nM) and ATP (2.5 mM) were mixed with the indicated concentrations of ^V13P^titin-BQ-ssrA at time zero to initiate binding. (B) Same as panel A but using the ^UF^titin-BQ-ssrA substrate. (C) Steady-state binding of ^UF^titin-BQ-ssrA to ^FS^ClpX (20 nM) in the presence of ATP (5 mM) or ATPγS (1 mM). (D) Changes in fluorescence during ^FS^ClpXP (10 nM) degradation of ^UF^titin-BQ-ssrA (0.1 or 20 μM) in the presence of ATPγS (0.5 mM). (E) Model for binding, engagement, and processive translocation/degradation. The transition from one translocation complex to the next (e.g., TC1 to TC2) is irreversible and involves a power stroke driven by ATP or ATPγS hydrolysis. Approximately 20 translocation steps would be required to fully translocate and degrade the titin substrate (5–9).

One prediction of our model is that sub-saturating concentrations of an unfolded substrate should show reduced steady-state binding to ClpX as a consequence of substrate release via processive translocation. Moreover, this difference should be greater in the presence of ATP, which supports faster translocation, than with ATPγS. Indeed, in the presence of ATP, ^FS^ClpX bound ^UF^titin-BQ-ssrA with an apparent affinity ~5-fold weaker (550 nM; Fig. 6C) than it bound titin-BQ-ssrA ATPγS (110 nM; Fig. 2B). In the presence of ATPγS, by contrast, the affinity constants for the native (59 nM) and unfolded substrates (90 nM) were stronger and differed by only ~1.5-fold. When we attempted to perform comparable ^UF^titin-BQ-ssrA binding experiments using ^FS^ClpXP, steady-state binding was not achieved as a consequence of rapid substrate degradation.

To determine how impaired hydrolysis influences binding, engagement, processive translocation, and degradation of the unfolded substrate, we conducted stopped flow association experiments with titin^UF^-BQ-ssrA and ^FS^ClpXP in the presence of ATPγS (Fig. 6D). Surprisingly, at high substrate concentrations, we observed oscillating fluorescence trajectories with a damped trend. At the highest substrate concentration, for example, the fluorescence initially decreased, then increased almost to the initial value, before decreasing, increasing, and decreasing again, with an approximate periodicity of 200 s. This time corresponds roughly to the time required to bind, translocate, and degrade initially bound substrate, and then begin this process anew but in a less synchronous manner. Oscillations are rare in biochemistry and require unusual kinetic mechanisms. One class of kinetic mechanisms that supports population coherence, defined as many molecules in the same states at the same time, involves numerous irreversible steps, each of which occur on a time scale that is short compared to the time of the overall reaction. In our experiment, a plausible explanation for the oscillating fluorescence is that there are two sets of states with distinct fluorescence intensities, which can only interconvert in a multi-step process. A likely possibility is that these states reflect initially bound and quenched states and a series of brighter translocating states, separated from each other by power strokes driven by irreversible ATPγS hydrolysis (Fig. 6E). By this model, the short steps are likely to reflect translocation of titin^UF^-BQ-ssrA through the axial channel of ClpXP.

## Discussion

We developed a fluorescence quenching assay to monitor binding of ssrA-tagged substrates to ClpX or ClpXP, revealing distinct hydrolysis-dependent kinetic processes between the initial substrate-binding event and the onset of unfolding, processive translocation, and degradation. The early substrate binding and engagement steps are fast, reversible, and convert an initial metastable recognition complex into a more stable complex poised for substrate unfolding and subsequent processive translocation and degradation.

In titration experiments, the apparent affinity constant (*K*_app_) of titin-BQ-ssrA for ^FS^ClpX varied from ~30-100 nM, with slightly stronger binding observed in the presence of ClpP, indicating that ClpP binding stabilizes a ClpX conformation with higher substrate affinity. In previous studies, *K*_M_ for ClpXP degradation of titin-ssrA, by contrast, was ~0.5-1.5 μM (18, 25). We suspect that the negatively charged and poly-aromatic BQ quencher may stabilize binding by making additional interaction with ClpX. Intriguingly, titin-BQ-ssrA quenched ^FS^ClpX fluorescence to a greater extent in the presence of ATPγS than ATP. Because ClpX and ClpXP hydrolyze ATPγS more slowly than ATP (21), this result suggested that the rate of hydrolysis alters the distribution of substrate-bound conformations. In support of this model, maximum quenching for titin-AQ-ssrA binding to ^FS^ClpXP was similar in the presence of ATP and ATPγS (Fig. 3C), as expected because a larger Förster radius makes quenching for this enzyme-substrate pair more efficient over longer fluorophore-quencher distances.

In kinetic experiments, we used the quenching assay to study substrate binding and engagement by ClpXP. The complex polyphasic fluorescence trajectories observed in association experiments using the titin-BQ-ssrA substrate (Figs. 3A and 3B) could not be fit to a model with fewer than three pre-unfolding enzyme-substrate complexes, each with their own characteristic mean fluorescence intensity. By contrast, titin-AQ-ssrA association trajectories were monophasic (Fig. 3C), supporting the idea that the distinct phases observed in the titin-BQ-ssrA association experiments represent distinct conformational states/ensembles with different average distances and/or orientations between the fluorophore on ClpX and the quencher on titin-BQ-ssrA.

A model including just the binding, intermediate, and engaged complexes provided excellent fits of the polyphasic association kinetics over a 200-fold range of titin-BQ-ssrA concentrations in the presence of ATP (Fig. 3A). As in any multistep kinetic modeling scheme, more complicated models could fit the results equally well or the same model with different rate constants might also provide adequate fits. Notably, however, the fitted rate constants and fluorescence intensities also accounted for dissociation kinetics (Fig. 4) and binding results (Fig. 2A; experimental and simulated *K*_app_ ≈ 60 nM). Based on the fitted fluorescence intensities, the quencher on the substrate appears to be closer to the TAMRA dye on ClpX in the intermediate complex than it is in the binding complex. This relationship makes sense if the quencher is farther above the axial channel in the binding than the intermediate complex (Fig. 5). We are leery of interpreting the fluorescence of the engaged complex in terms of quencher-fluorophore distance alone, as the microenvironment of the quencher and/or fluorophore in this state seem likely to also affect energy transfer via orientation effects. For example, the negatively charged quencher is expected to be in or near the top of the ClpX axial channel, where it could interact with positively charged ClpX RKH loops, and the fluorophore on the top face of ClpX may contact the native titin domain.

Both titin-BQ-ssrA association and dissociation were slower in the presence of ATPγS than ATP, and the fitted values of the *k*_2_, *k*_-2_, *k*_3_, and *k*_-3_ rate constants from the ATPγS experiments were 3-7 fold smaller than those determined in the ATP experiments (Table 1). These results suggest that the rate of ATP/ATPγS hydrolysis is an important factor in determining the rates of the associated conformational transitions. Under the conditions of our experiments, the maximum steady-state rate of ATP hydrolysis by ClpXP is ~3.6 s^−1^ (28). The fitted values of the *k*_2_, *k*_-2_, *k*_3_, and *k*_-3_ rate constants in experiments performed using ATP were all smaller than this hydrolysis rate (Table 1), and thus some of these steps might include multiple hydrolysis reactions.

The number of states uniquely resolvable in our stopped-flow experiments limits the complexity of our model. Moreover, kinetic states in the model may not be unique conformations but rather ensembles of conformations explored in a semi-synchronous manner upon rapid mixing. For example, in our fits, the fluorescence value of the engaged complex is lower in ATPγS than in ATP. Thus, it is likely that this complex represents an ensemble average of low fluorescence pre-ATP hydrolysis states (more highly populated in ATPγS) and higher fluorescence post-ATP hydrolysis states (relatively more populated in ATP).

In experiments with natively destabilized or unfolded titin substrates, we explored the kinetic relationships between binding, engagement, and degradation. Binding and engagement of the destabilized ^V13P^titin-BQ-ssrA substrate by ^FS^ClpXP resembled that of titin-BQ-ssrA. At high ^V13P^titin-BQ-ssrA concentrations, transitions relating to binding and engagement were largely complete with 5 s of mixing, whereas unfolding, processive translocation, and degradation were substantially slower. Association experiments with ^UF^titin-BQ-ssrA and ^FS^ClpXP also revealed rapid association and engagement, relative to degradation. The intermediate steady-state fluorescence observed at late time points with high ^UF^titin-BQ-ssrA concentrations presumably reflects a mixture of different ClpXP-substrate complexes, distinct from the ensemble of states observed with titin-BQ-ssrA (prior to unfolding), and thus is refractory to direct comparison.

Specific degron recognition by ClpXP and related AAA+ proteases is necessary to avoid uncontrolled protein destruction, as later steps including substrate unfolding and processive translocation through the axial channel have little specificity (18–19; 33). The specificity of recognition and promiscuity of processive translocation seem at first to be at odds. A highly specific AAA+ protease might be expected to bind tightly to a limited set of sequences, potentially limiting processivity, whereas a highly processive protease that can grip most sequences for translocation might be expected to have limited specificity. As outlined below, the kinetic studies presented here together with recent cryo-EM structures of substrate-bound ClpXP (28–30) suggest a mechanism by which ClpXP achieves specificity and processivity.

Our studies provide evidence for three sequentially populated substrate-bound ClpXP complexes (BC, IC, and EC) prior to substrate unfolding (Fig. 5). For the titin-BQ-ssrA substrate, the relative pseudo-equilibrium populations of BC, IC, and EC are approximately 13%, 8%, and 79%, respectively. Thus, IC is a higher energy state than BC, and EC is the lowest energy state. BC is likely to be similar to a cryo-EM structure in which an ssrA degron binds in the top portion of an otherwise closed ClpX channel with specific packing and hydrogen-bond interactions between ClpX side chains and the Ala-Ala-COO^−^ of the ssrA tag (29). IC appears to correspond to a cryo-EM structure in which the ssrA degron moves six residues deeper into a now open ClpX channel, but the native portion of the substrate is not in contact with the top of the ClpX ring (29). EC appears to be similar to a cryo-EM structure in which the degron has moved deeply enough into the channel to draw the native portion of the substrate against the entrance to the ClpX channel (28). Substrate contacts in the channels of the IC-like and EC-like structures appear to be generally similar to each other and non-specific (28, 29). Hence, the lower energy of EC relative to IC is likely to result from contacts between the native portion of the substrate and ClpX.

The BC-like conformation of ClpXP is ideally suited for ‘checking’ short unstructured peptides in proteins for degrons, as the contacts with the Ala-Ala-COO^−^ of the ssrA-degron in this complex are sequence dependent (29). Although contacts in the downstream IC and EC complexes are non-specific, our results indicate that both of these structures can revert to BC. For example, once titin-BQ-ssrA reaches IC, it is ~4-fold more likely to revert back to BC than to move forward to EC. Even though the ssrA degron has evolved to ensure rapid ClpXP degradation of attached proteins, only ~20% of titin-BQ-ssrA substrates that bind initially to form BC advance directly to EC, largely as a consequence of this back step. Because this IC→BC step is ATP dependent, this cycle could potentially act as a type of kinetic proofreading (34) by providing multiple opportunities for substrate discrimination. Indeed, the multiple ATP-hydrolysis-dependent steps that we observe prior to substrate engagement should ensure that only substrates with proper degrons have a substantial chance of being unfolded and degraded. Thus, degron interactions in the BC-like structure appear to mediate ClpXP specificity, whereas ATP-dependent translocation of the degron deeper into the axial channel allows ClpXP to reversibly transition into a more promiscuous and ultimately processive unfoldase/translocase. We note that the detailed mechanism of substrate engagement by ClpXP is likely to depend upon degron length (29), and it will be important to determine how changes in tag length and sequence affect mechanism using the methods developed here. It is possible that other AAA+ proteases use similar multi-step engagement mechanisms to ensure specific and robust destruction of their targets.

## Methods

### Proteins

All proteins contained His_6_ tags, were expressed from plasmids in *E. coli*, and were purified using established procedures (35, 36). A single-chain *E. coli* ^C169S^ClpX^ΔN^ pseudohexamer with a single solvent-exposed Cys^170^ residue in subunit 1 was constructed by sortase-mediated ligation of appropriate single-chain trimers as described (36). This pseudohexamer was labeled with TAMRA-maleimide (ThermoFisher Scientific) to produce ^FS^ClpX. BHQ-10 maleimide was prepared from BHQ-10 succinimidyl ester (Biosearch Technologies) as described for the related BHQ-3 molecule (36). Our ssrA-tagged titin I27 construct contained the N-terminal sequence Met-His-Glu-Gly, residues 1-89 of the I27 domain of human titin (NCBI accession 1530732605 or 6I0Y_z), followed by the sequence Gly-Cys-Gly-(His)_6_-<u>Ala-Ala-Asn-Asp-Glu-Asn-Tyr-Ala-Leu-Ala-Ala</u> (ssrA tag underlined). To produce titin-BQ-ssrA, this protein was labeled for 30 min at room temperature with three molar equivalents of BHQ-10 maleimide in buffer A (20 mM HEPES (pH 7.7), 30 mM KCl, 0.1 mM EDTA, 10% glycerol) supplemented with 5 mM Tris (pH 8.0). Following labeling, the protein was desalted into buffer A using a PD10 column, and titin-BQ-ssrA was separated from unlabeled protein by SOURCE-15Q chromatography using a gradient from 0-40% buffer B (20 mM HEPES (pH 7.7), 1 M KCl, 0.1 mM EDTA, 10% glycerol). Fractions containing pure titin-BQ-ssrA were pooled, concentrated, and flash frozen. To produce titin-AQ-ssrA, our ssrA-tagged titin protein was reacted at room temperature with three equivalents of ATTO 575Q maleimide (ATTO-TEC; GMBH) for 30 min in buffer A supplemented with 5 mM Tris (pH 8.0). The reaction was quenched by addition of 15 mM DTT, desalted using a PD-10 column, desalted further by Sephadex G-25 chromatography, and then concentrated and flash frozen. The concentrations of titin-BQ-ssrA and titin-AQ-ssrA were measured by the BCA assay (ThermoFisher Scientific).

### Binding and kinetic assays

Equilibrium and kinetic experiments as well as degradation assays were performed at room temperature in PD buffer (25 mM HEPES (pH 7.5), 100 mM KCl, 10 mM MgCl_2_, 0.1 mM EDTA, 10% glycerol) supplemented with ATP (5 mM) or ATPγS (1 mM) as indicated. Experiments performed with ATP also contained a creatine/creatine-kinase regeneration system (16 mM phosphocreatine and 0.032 mg/mL creatine kinase; Sigma). For equilibrium binding, fixed concentrations of ^FS^ClpX (typically 20 nM) without or with ClpP (60 nM) were initially mixed in the absence of nucleotide for 2 min with different concentrations of titin-BQ-ssrA or titin-AQ-ssrA in a microtiter plate. Under these conditions, ^FS^ClpX does not bind substrate. Fluorescence emission spectra from 560-590 nm (excitation 530 nm) were recorded using a SpectraMax M5 plate reader (Molecular Devices). Next, ATP or ATPγS was added. After incubation for 5 min (ATP experiments) or 15 min (ATPγS experiments), a second fluorescence emission spectrum was recorded. Areas under emission curves were calculated without nucleotide (F_buf_) and with nucleotide (F_nuc_) and the equilibrium fluorescence quenching value was defined as 1 – F_nuc_/F_buf_ – Q_buf_, where Q_buf_ was the slight increase in fluorescence that occurred with dilution of the nucleotide-free control.

Kinetic experiments were performed using an SF-300X stopped-flow instrument (KinTek). For association kinetics, one syringe contained a fixed concentration of ^FS^ClpX, ClpP, ATP or ATPγS, and a second syringe contained varying concentrations of titin-BQ-ssrA or titin-AQ-ssrA. For dissociation experiments, titin-BQ-ssrA was incubated for 15 min with ^FS^ClpX, ClpP, and ATP or ATPγS and loaded into one syringe. The second syringe contained 50 μM unlabeled I27 titin-ssrA. Following mixing of the syringe contents (dead time ~1 ms), the fluorescence trajectory was recorded (excitation 550 nm; emission passed through a Newport 580 nm/10 nm band-pass filter).

Stopped-flow kinetic data was globally fit using KinTek Explorer software (KinTek). The general model was developed from the ClpXP and titin-BQ-ssrA association experiments conducted in ATP (Fig. 3A). The fluorescence traces were fit to models with one, two, three, and four bound states, each with distinct fluorescence intensities. For simplicity, strictly linear progression or retrogression between states was enforced. Forward and reverse rate constants and state-specific fluorescent intensities were allowed to vary freely during the fitting process, and the parameter space was comprehensively explored. We were not able to obtain satisfactory fits with models that included fewer than three bound states.

We fit the data from the ClpXP/titin-BQ-ssrA/ATPγS association experiments (Fig. 3B) to the same model with three substrate bound states. To obtain satisfactory fits, we needed to vary not just the rate constants controlling the interchange between states but also the state-specific fluorescent intensities, suggesting that the ensembles of conformations reflected by each bound state differ between ATP and ATPγS.

We did not include dissociation data in the fitting process, but rather confirmed that our fits based on association data were also consistent with dissociation data. From the association-derived fits, we calculated the distribution of states at steady state with quencher-labeled substrate and simulated the changes in fluorescence intensity upon introduction of a vast excess of unlabeled substrate. For both ATP and ATPγS, we found that, the simulated fluorescence changes were very similar to those observed in dissociation experiments.

Satisfactory fits of data from ^V13P^titin-BQ-ssrA and ^UF^titin-BQ-ssrA association/degradation experiments (Figs. 6A and 6B, respectively) required an additional bound state and an irreversible proteolytic step. We maintained the rates and fluorescent intensities for the bimolecular association and the interconversion between the first three bound states as close to those of the titin-BQ-ssrA model as possible while maintaining a satisfactory fit. Models with many parameters have many possible solutions, and thus our goal was to find example parameter combinations that were broadly consistent with the experimental data.

